# reactifpTM: an accessible reimplementation of actifpTM

**DOI:** 10.64898/2026.08.24.746624

**Authors:** Adam J. Simpkin, Emily J. Johnson, Daniel J. Rigden

## Abstract

**Motivation:** The actual interface pTM score (actifpTM) is a modified version of the ipTM score that limits the calculation to only those residues at the interface. Whilst actifpTM provides an effective interface quality score, a limiting factor is that it makes use of the predicted aligned error (PAE) with probabilities, information that is generated during a ColabFold run, but not output by the package or other model prediction software. The consequent inability to generate actifpTM scores for the results of software such as AlphaFold 2 or AlphaFold 3 has limited its adoption. With reactifpTM we address this problem by providing a standalone tool that can be run on the standard outputs of most model prediction packages.

**Results:** Using the same underlying principles as actifpTM, reactifpTM has been developed to use standard output files from model prediction software (a model and corresponding PAE) to perform an actifpTM-like calculation. ColabFold models were generated for a dataset of 1079 known interfaces in the PDB. A strong correlation was shown between actifpTM and reactifpTM for this dataset.

**Availability and implementation:** reactifpTM is coded in Python. All scripts and associated documentation are available from https://github.com/hlasimpk/reactifptm or https://pypi.org/project/reactifptm.

## 1. Introduction

AlphaFold (Jumper et al. 2021; Abramson et al. 2024) and other model prediction tools (Mirdita et al. 2022; Wohlwend et al. 2025; Passaro et al. 2025; Chai Discovery et al. 2024; The OpenFold3 Team 2025; ByteDance AML AI4Science Team et al. 2025; Corley et al. 2025) have had a profound impact on the field of structural biology, allowing the easy generation of high quality models of proteins and complexes, alongside effective scoring metrics to assess model quality. These include the interface predicted template modeling score (ipTM), introduced with AlphaFold-multimer (Evans et al. 2021) to evaluate the confidence of predicted binding interfaces. The availability of these tools has enabled large-scale *in silico* protein-protein interaction (PPIs) screens (eg Schmid et al. 2025), with the AlphaFold Database (AFDB) now extended to include homo- and heterodimeric complexes (Varadi et al. 2022; Fleming et al. 2025). There is therefore an increasing demand for metrics that can distinguish true PPIs from false positives.

Whilst ipTM is an effective score in many circumstances, often as part of a weighted average with the per-chain pTM scores (Evans et al. 2021), several studies have found that varying the lengths of interacting sequences resulted in differing ipTM scores despite identical binding mechanisms (Lee et al. 2024; Bret et al. 2024). Specifically, longer sequences containing disordered regions tend to yield lower ipTM scores, as disordered residues are weighted equally in the calculation despite not contributing to the interface. An obvious solution would be to only use truncated sequences excluding disordered regions, but this has been shown to have a deleterious effect on the models (Lee et al. 2024; Bret et al. 2024).

To counter this problem, several metrics have been developed to use the PAE including the actifpTM score (Varga et al. 2025) and the ipSAE score (Dunbrack 2025). Both of these scores have been shown to be agnostic to variations in sequence length. The actifpTM score is available within ColabFold and takes internal parameters as inputs, whereas ipSAE is a standalone package that uses the common output files from AlphaFold 2 and 3. This broader range of usage scenarios has undoubtedly contributed to the rapid adoption of the ipSAE score in cases as diverse as host-pathogen studies (Zhang 2026) to metabolon characterisation (Yu King Hing et al. 2026).

Here we introduce reactifpTM, a reimplementation of the actifpTM score designed to operate only on widely available outputs from the latest generation of model prediction software and, thereby, to have broad applicability. We show that the score correlates closely with original actifpTM on a dataset of 1079 complexes, and that it can complement the ipSAE score.

## 2. Methods

### 2.1. reactifpTM methodology

The actifpTM score primarily differs from ipTM in the residue weights set for each interchain residue pair. In the ipTM score, the weights are all set to 1, whereas in the actifpTM score, weights vary in a structure-dependent fashion. The contact probability map is used to calculate the probability that each interchain residue pair is within 8 Å of one-another (Cβ-Cβ atoms, C□ for glycines) producing a pseudo-contact map. Using these contact probabilities as weights effectively masks out residue pairs which are far apart in space, and therefore allows the actifpTM to only focus on the residue pairs at the interface of the complex.

Where contact probabilities are output (e.g. AlphaFold 3) this strategy could be replicated. More often, however, contact probabilities are not output by model prediction software and therefore another strategy is required to establish a more general score. For this we use the output models directly. The Euclidean distance between each residue pair is calculated (Cβ-Cβ atoms, C□ for glycines) and a contact map is created from residue pairs found to be within 8 Å of one-another. As before, this contact map is used to mask out residue pairs that are distant from one-another, with remaining pairs assigned a weight of 1.

After adjusting the residue weights, actifpTM essentially performs a standard ipTM calculation where the pairwise pTM-score matrix is calculated from the PAE probability matrix using:

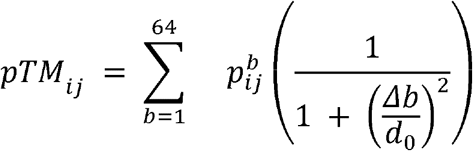

Since the PAE matrices (n, n) output by software such as AF3 are derived from the PAE probability matrix (n, n, 64) using:

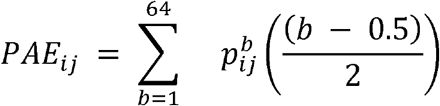

We are unable to calculate the *pTM*_*ij*_ value directly. However, the ipSAE score introduced an approximation which is also adopted here:

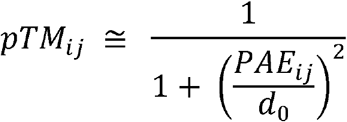

When confidence is high and the probability is concentrated in a narrow range of bins, the two expressions are nearly equivalent. The approximation therefore performs best for high-confidence interfaces that are most relevant to interaction prediction.

### 2.2. Examples from actifpTM and ipSAE

To validate reactifpTM against existing metrics, models were generated for six test cases drawn from the actifpTM and ipSAE publications using ColabFold v1.6.1 with the --calc-extra-ptm flag. Full details of these cases are provided in Supplementary Table 1.

### 2.3. DockGround dataset generation

A set of complexes was downloaded from the DockGround database (Collins et al. 2022). Dockground’s filtering system was used to limit the dataset to only include heterodimeric protein-protein complexes published in the PDB between 19/05/1976 and 17/04/2026 with ≤3 Å resolution, followed by redundancy reduction using 30% sequence identity. The dataset was further filtered manually to remove antibody/antigen complexes and complexes containing more than 1550 amino acids in total due to modelling constraints. This left a total of 1079 complexes.

ColabFold (Mirdita et al. 2022) version 1.6.1 was used to generate 5 models for each of the complexes. The --calc-extra-ptm flag was set to generate the actifpTM score for each model.

## 3. Results

### 3.1. Examples from actifpTM and ipSAE

Across the six validation cases drawn from the actifpTM and ipSAE publications (Supplementary Table 1), reactifpTM closely reproduced actifpTM scores, including the key behaviour of maintaining consistent scores for both short and long peptide forms of the same interaction. Notably, ipSAE gave low scores for two of the short peptide cases (SH3BP5/MAPK and KEAP1/NF2L2). We confirmed that both peptides were correctly placed in the binding site, suggesting that ipSAE may be less robust when applied to peptide interactions with limited interchain contacts. The two negative control cases from the ipSAE paper returned low scores for both metrics, with reactifpTM again closely matching actifpTM in both cases.

### 3.2 DockGround dataset results

Figure 1 shows a larger scale comparison of reactifpTM with actifpTM and ipSAE. The reactifpTM correlates well with the actifpTM (Fig 1A, Pearson’s r: 0.990, Spearman’s ρ: 0.991), but it is clear that the actifpTM often scores higher than the reactifpTM, especially for low-confidence interfaces (actifpTM score <0.8). This can be seen more clearly when plotted as a Bland-Altman plot (Suppl. Fig 1) where actifpTM - reactifpTM is plotted against the mean of both scores. We can see that above 0.8, the difference is often close to zero, but below 0.8 the difference tends to be positive.

**Figure 1.**
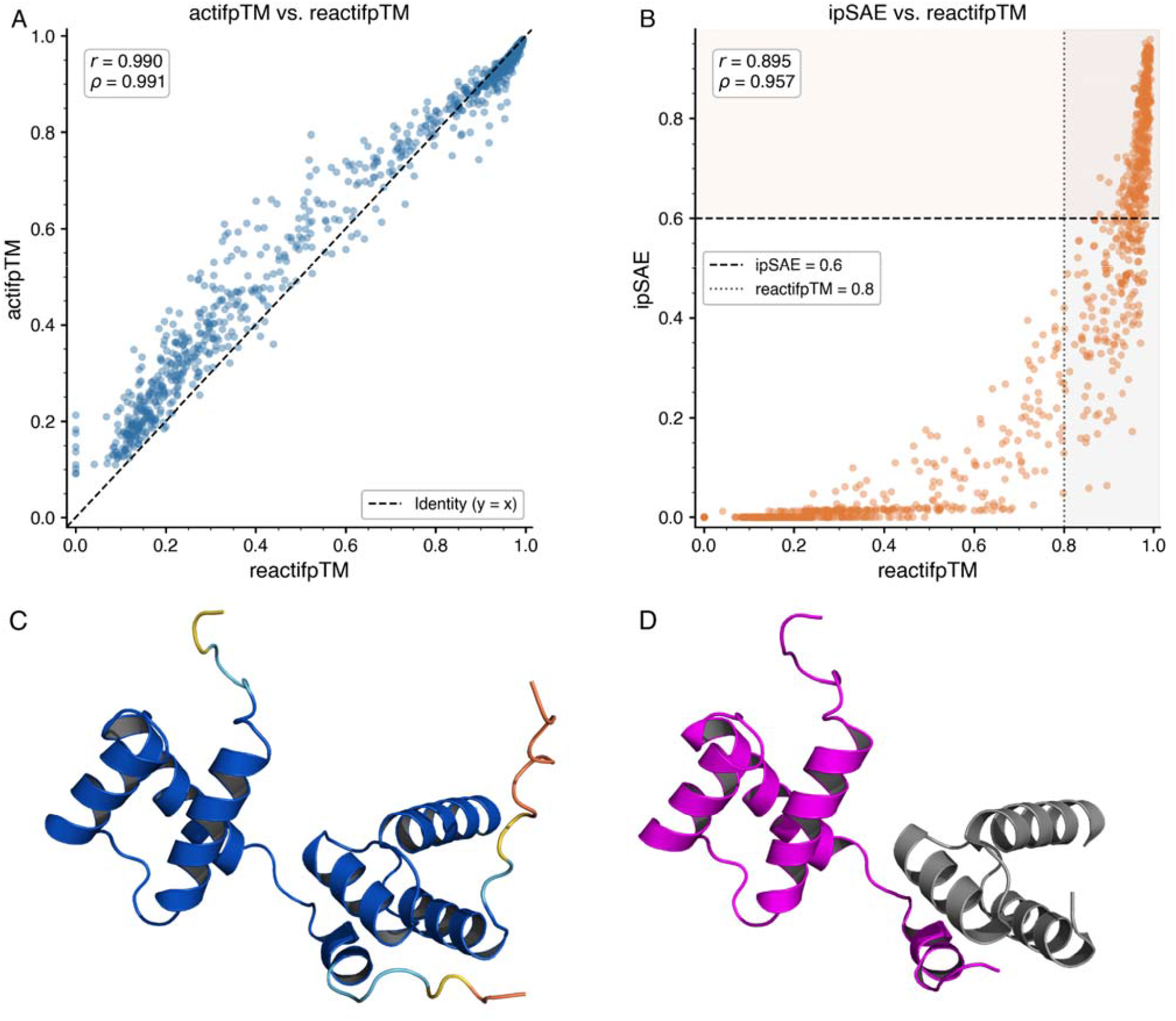
A) The comparison of actifpTM and reactifpTM on the DockGround derived dataset. The Pearson’s r correlation score and a line indicating perfect correlation are shown. B) The comparison of ipSAE and reactifpTM on the DockGround derived dataset. The Pearson’s r correlation coefficient, Spearman’s ρ correlation coefficient, and threshold indicating a good interface are shown for ipSAE and reactifpTM are shown. C) ColabFold model of 1AKH (chains A & B) coloured by pLDDT on a blue to orange scale where blue signifies high pLDDT D) The crystal structure of 1AKH where chain A is shown in grey and chain B is shown in magenta.

This can be explained by the way that the actifpTM and reactifpTM calculate *pTM*_*ij*_. For high-confidence interfaces, the probabilities are narrowly distributed, meaning the approximate and actual *pTM*_*ij*_ values are nearly equivalent. For lower confidence interfaces, the distribution broadens across bins, causing the approximate *pTM*_*ij*_ to deviate from the actual *pTM*_*ij*_. The effect of this is shown clearly in Supplementary Figure 2 where we explore 9G25_H_K, a case where the actifpTM was significantly higher than the reactifpTM (0.658 vs 0.401).

We also noted that in some scenarios, reactifpTM scored zero whilst actifpTM gave scores between 0.093 and 0.213. This occurs when no interchain pairs are found by reactifpTM, i.e. when the distances between chains are greater than 8 Å (Suppl. Fig 3). The distogram method used by actifpTM does not rely on the final coordinates. Therefore actifpTM can assign non-zero interface scores when the distogram encodes a weak preference for proximity that is not realised in the relaxed structure. In such cases, reactifpTM’s coordinate-based contact criterion may be preferable as it only scores interfaces that are actually present in the output model.

The reactifpTM score also correlates well with the ipSAE score (Fig 1B, Pearson’s r: 0.895, Spearman’s ρ: 0.957), although we can see a number of cases where reactifpTM is confident in the interface and ipSAE is not. Fig 1C/D shows example 1AKH_A_B, which had confident actifpTM and reactifpTM scores (0.931 and 0.932 respectively), but a much less confident ipSAE score (0.54). The TM score (Zhang and Skolnick 2004) to the crystal structure was 0.79, indicating the model had accurately captured the interface. This pattern was most frequently observed in cases with fewer interchain residue contacts; for example, ipSAE identified only 82 interchain pairs for 1AKH_A_B.

## 4. Discussion

The ability to reliably score predicted model complexes has become increasingly important as model prediction software has allowed large scale *in silico PPI screeni*ng of proteomes across model organisms (Schmid et al. 2025; Kim et al. 2026; Yu et al. 2023). The incorporation of predicted complexes in the AFDB has further amplified this need. When assessing models at this scale, even modest differences between metrics can translate to thousands of false negative or false positive interaction predictions.

Here, we introduce the reactifpTM as a more generally accessible proxy for the well-known actifpTM score. The two scores correspond well, especially for high-confidence interfaces. A key difference is that reactifpTM will often give lower scores for lower confidence interfaces as a result of not using the PAE with probabilities. We consider this a reasonable tradeoff in order to allow the score to be calculated for the results of more model prediction packages.

A variety of other metrics have been introduced to for *in silico* PPI screening including pDockQ/pDockQ2 (Bryant et al. 2022; Zhu et al. 2023), LIS (Kim et al. 2024), iLIS (Kim et al. 2026), each capturing subtly different aspects of interface confidence. Among these, ipSAE has seen rapid and widespread adoption as a PAE-based interface scoring metric, making it a natural point of comparison for validating reactifpTM as a complementary approach addressing the same class of problem. Here we focused on validating reactifpTM and demonstrating that it reproduces actifpTM scores accurately across a benchmark of known structures. In doing so, we also highlight cases where both actifpTM and reactifpTM identify confident interfaces that fall below the ipSAE threshold, as illustrated by 9MT6_B_C, where ipSAE’s d0 scaling to interface size appears to penalise small but genuinely confident interfaces. The assumptions embedded in the design of each metric may favour particular complex types, and further work is needed to establish their relative strengths and limitations. We would therefore recommend using reactifpTM alongside ipSAE as complementary tools, particularly where interfaces involve short peptides or limited interchain contacts.

## 5. Figures and supplementary

**Supplementary Table 1).**
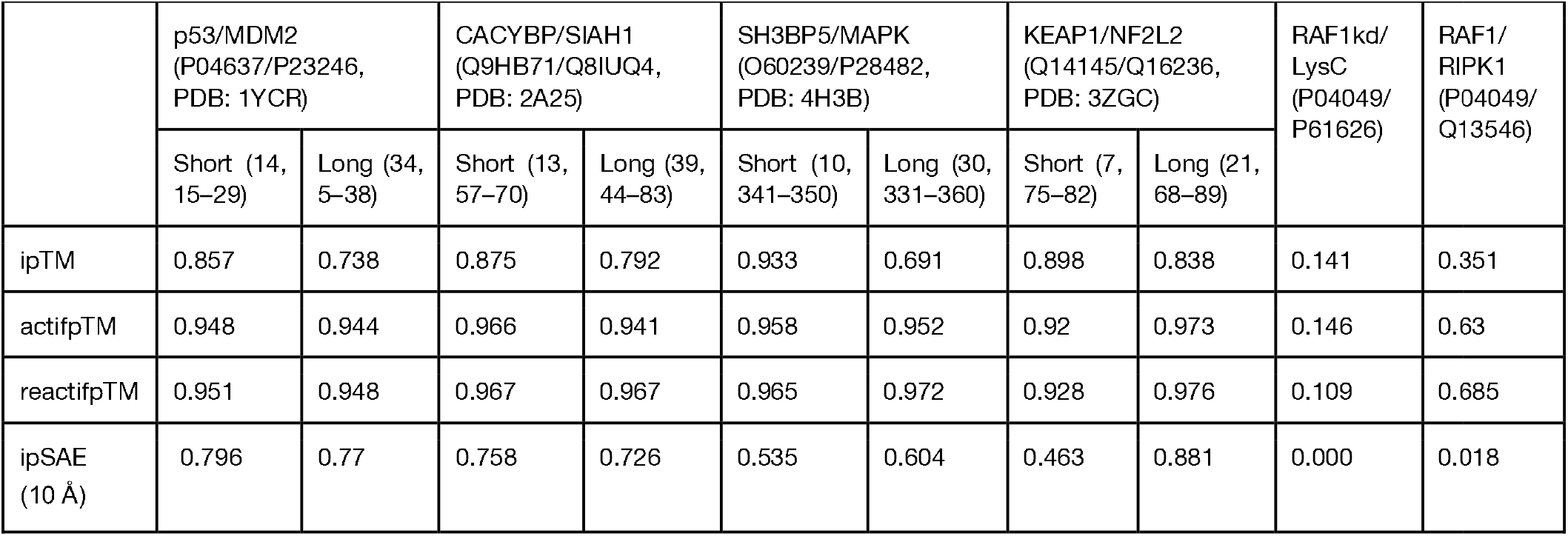
Comparison of ipTM, actifpTM, reactifpTM and ipSAE scores for validation cases drawn from the actifpTM and ipSAE publications. UniProt accession codes and PDB codes (where available) are given in parentheses alongside each complex. The first four cases are taken from the actifpTM paper; Short and Long distinguish scores for short peptides of PDB-deposited length and extended versions containing flanking regions of equal length (peptide lengths and residue boundaries shown in parentheses, UniProt numbering). The final two cases are taken from the ipSAE paper as negative controls, where RAF1kd represents the kinase domain of RAF1 (residues 17–289) and RAF1, LysC and RIPK1 are full-length UniProt sequences. All models were generated with ColabFold v1.6.1.

**Supplementary Figure 1.**
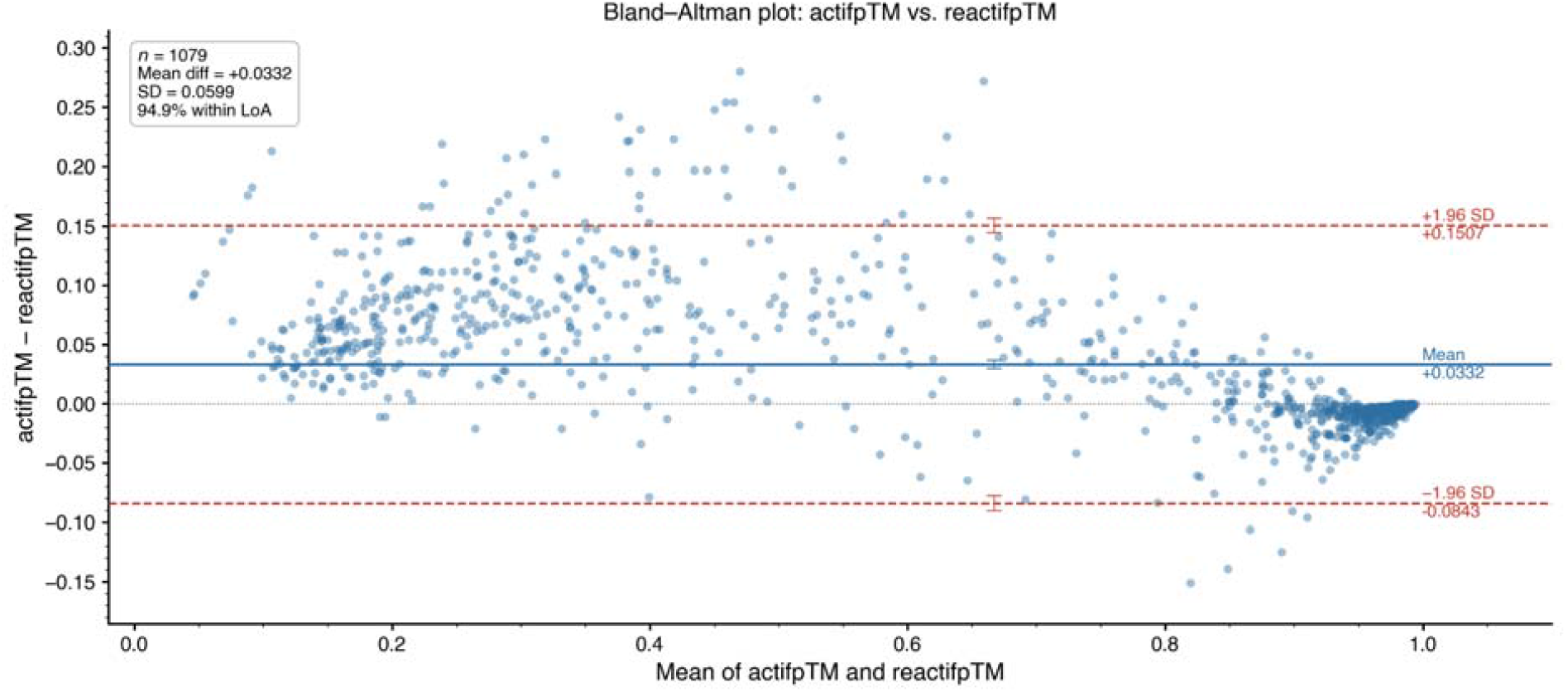
A Bland-Altman plot comparing actifpTM and reactifpTM scores across 1079 complexes. The y-axis shows the difference (actifpTM - reactifpTM) and the x-axis shows the mean of the two scores. The solid blue line shows the mean bias (0.0332), and the dashed red lines show the 95% limits of agreement (0.1507, - 0.0843). Below a mean score of 0.8, reactifpTM tends to score lower than actifpTM, reflecting the differences to *pTM*_*ij*_ caused by using the PAE without probabilities. Above 0.8, the two methods converge, suggesting high-confidence interfaces are robustly captured by both approaches.

**Supplementary Figure 2.**
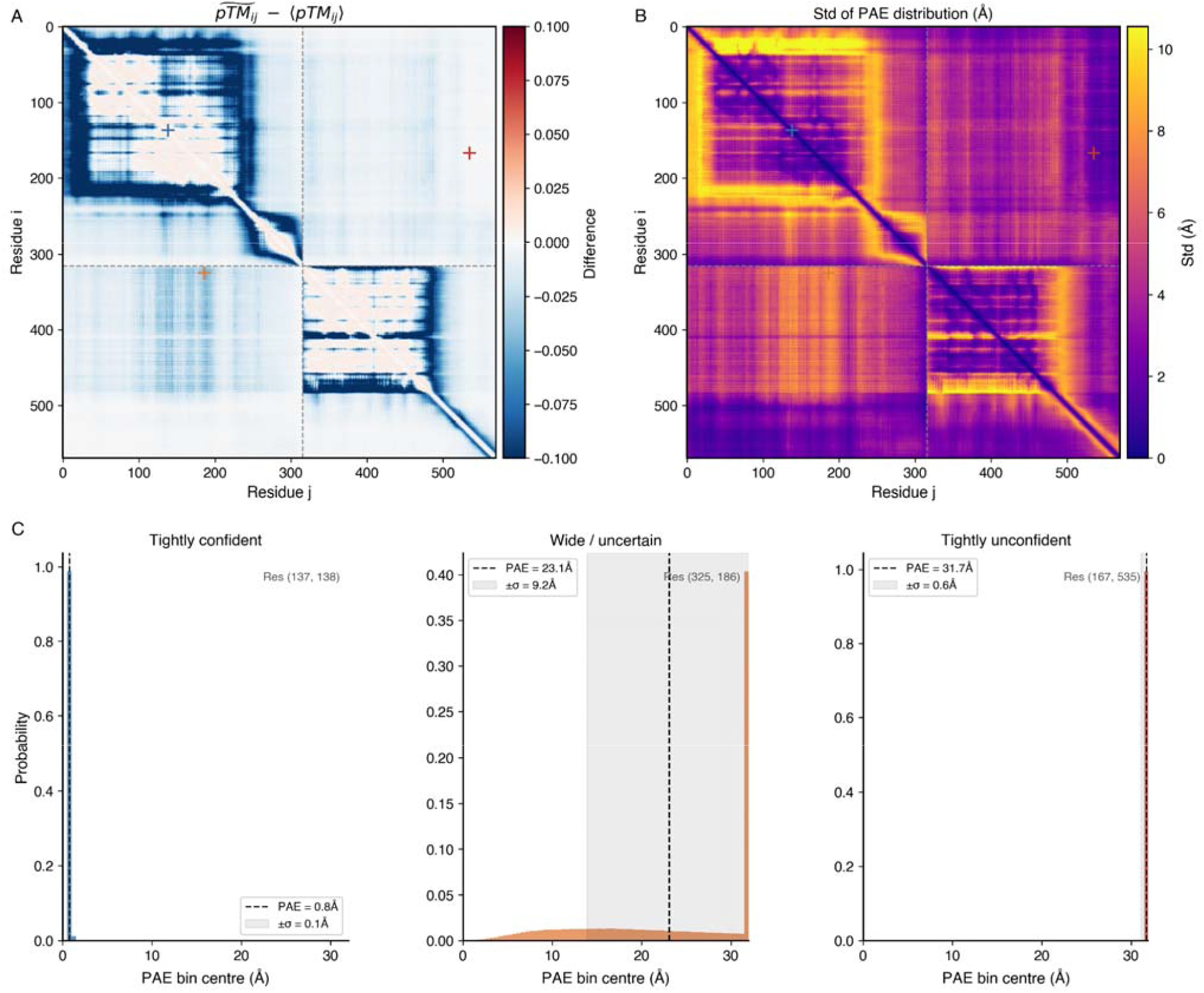
A comparison between the approximate *pTM*_*ij*_ score and the *pTM*_*ij*_ score for 9G25_H_K. A) shows the difference between the two scores, where dark blue regions highlight residue pairs where the approximate pTM_ij_ is less than the *pTM*_*ij*_. B) shows the standard deviation of the probability bins for each residue pair. We can see that the areas where the standard deviation is greatest correspond to the regions where the approximate *pTM*_*ij*_ deviates from the *pTM*_*ij*_. C) shows the probability distributions for tightly confident (sharp peak t low PAE), wide/ uncertain (broad distribution), and tightly unconfident (sharp peak at high PAE) residue pairs. It s the wide/uncertain PAE distributions that lead to differences between the approximate *pTM*_*ij*_ and the *pTM*_*ij*_.

**Supplementary Figure 3.**
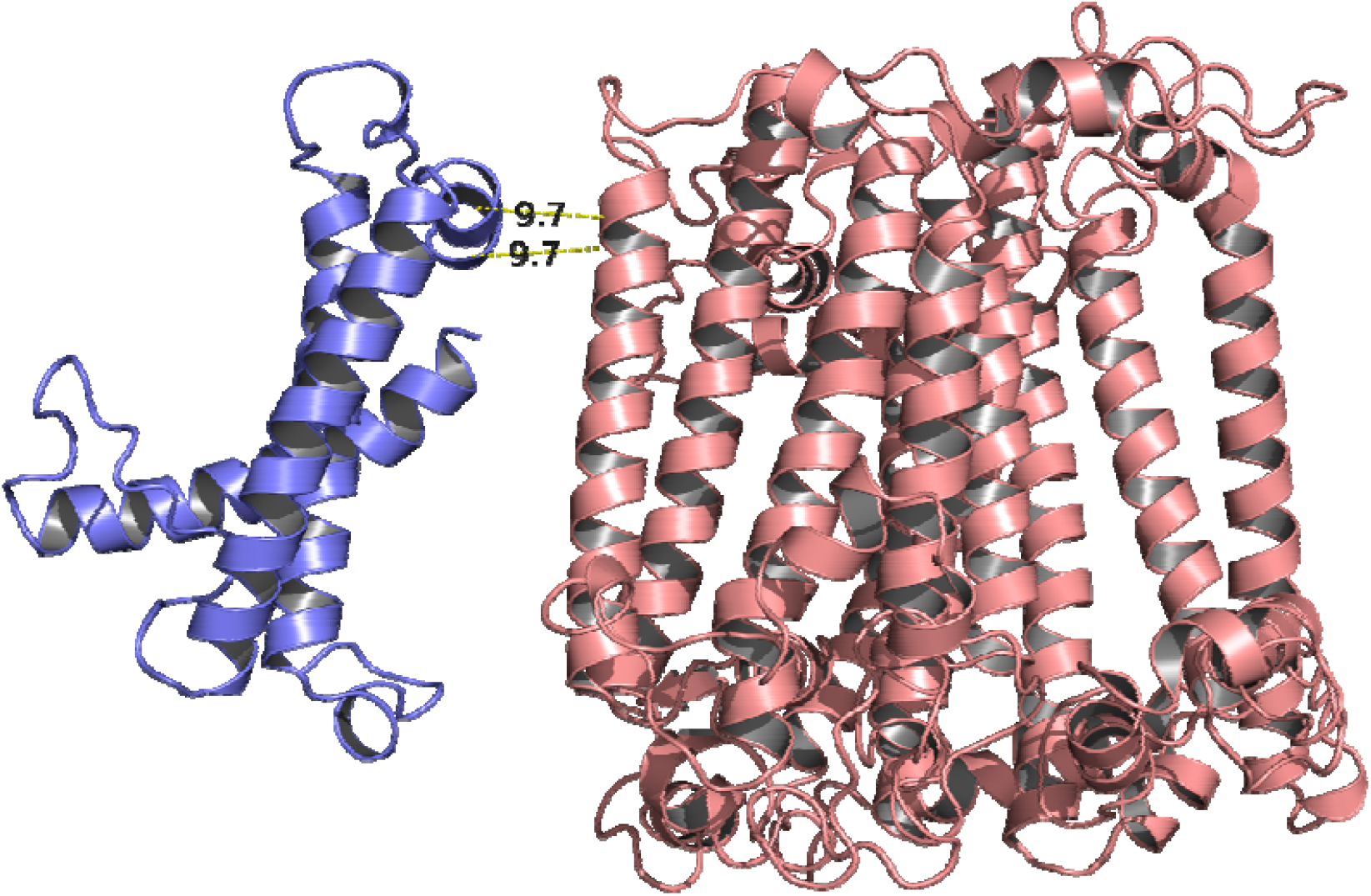
Visualisation of chain P (auth id 3; slate) and chain B (salmon) for 9KC5_3_B. This model gave a reactifpTM score of 0.0 and an actifpTM of 0.176. The minimum Cβ–Cβ distance between the 2 chains is 9.7 □, which explains why reactifpTM returns a zero.

